# Conformational changes in the AdeB transmembrane efflux pump by amphiphilic peptide Mastoparan-B, down-regulates expression of the *ade*B Gene and restores antibiotics Susceptibility

**DOI:** 10.1101/2023.01.03.522678

**Authors:** Mohammad Reza Shakibaie, Farzan Modaresi, Omid Azizi, Omid Tadjrobehkar, Mohammad Mehdi Ghaemi

**Affiliations:** Department of Microbiology and Virology, Kerman University of Medical Sciences, Kerman, Iran; Gastroenterology Hepatology Research Center, Institute of Basic and Clinical Physiology Sciences, Kerman University of Medical Sciences, Kerman, Iran; Department of Microbiology, School of Medicine, Jahrom University of Medical Sciences, Jahrom, Iran; Department of laboratory sciences, Torbat Heydariyeh University of Medical Sciences, Torbat Heydariyeh, Iran; Medical Informatics Research Center, Institute for Futures Studies in Health, Kerman University of Medical Sciences, Kerman, Iran

**Keywords:** Mastoparan-B, molecular docking, efflux process, protein modeling, AlphaFold2

## Abstract

No report exists on the role of Mastoparan B (MP-B) as an RND efflux pump inhibitor in multi-drug resistant (MDR) *Acinetobacter baumannii*. Here, we performed a series of in-silico experiments to predict the inhibition of the AdeB efflux pump by MP-B as a drug target agent. For this reason, an MDR strain of *A. baumannii* was subjected to MICs against 12 antibiotics as well as MP-B. Expression of the a*de*B gene in the presence and absence of sub-MIC of MP-B was studied by qRT-PCR. It was found that MP-B had potent antimicrobial activity (MIC=1 μg/ml) associated with a 20-fold decrease in the *ade*B gene expression at the sub-MIC level. The stereochemical analysis using several automated servers confirmed that the AdeB protein is an inner membrane of the RND tripartite complex system with helix-turn-helix conformation and a pore rich in Phe, Ala, and Lys residue. Furthermore, 20 ligands were generated from the initial docked poses to create the correct protein-peptide complexes using the BioLiP pipeline. The pose showed high Z=1.2, C=1.41, TM=0.99, and RMSD=4.4 scores was selected for docking purposes. The molecular docking via AutoDock/Vina revealed that MP-B form H-bound with Val 499, Phe 454, Thr 474, Ser 461, Gly 465, and Tyr 468 residues of the AdeB helix-5 and caused a shift in the dihedral angle (Φ/Ψ) by distances of 9.0 Å, 9.3 Å, and 9.6 Å, respectively. This shift in folding was detected by AlphaFold 2 and influenced the overall druggability of the protein. From the above results, we concluded that MP-B can be a good candidate for bacterial efflux pump inhibition.

---

**C**omputational chemistry has greatly increased our knowledge about protein conformation and drug interaction and helped us better understand microbial infection treatment (1). Biomolecular simulation with models allows for investigations of target proteins’ structure and function features, which are useful for identifying drug-binding sites and elucidating drug action mechanisms (2,3). Recently a breakthrough in the 3D construction of various proteins occurred with the invention of the AlphaFold 2 Protein Structure Database in cooperation with EMBL-EBI. AlphaFold 2 can accurately predict 3D models of protein structures and is accelerating research in nearly every field of biology such as drug–target interaction (4).

In this regard, *A. baumannii* is a Gram-negative hospital-acquired pathogen to blame for high morbidity and mortality within the medical care units such as ICUs of various hospitals across the world (5). Following the increasing emergence of strains that are not susceptible to any clinically used antibiotics, the WHO has ranked carbapenem-resistant strains of this bacterium in first place in its global priority pathogen list (6). *A. baumannii* illustrate different resistance mechanisms against antimicrobial agents such as β-Lactam inactivation, biofilm formation, efflux pumps, alterations of the affinity for antibiotics, etc (7, 8). One of the most important mechanisms of resistance to antibiotics is the efflux pump which plays a key role in the intrinsic resistance to several antibiotics (9-11). The efflux pumps are divided into five superfamilies: ATP-binding cassette (ABC), small multi-drug resistance (SMR), multi-antimicrobial extrusion (MATE), major facilitator (MFS), and resistance/nodulation/division (RND) (12). The best-characterized multi-drug efflux system in *A. baumannii* is the RND superfamily (13). These tripartite complexes comprise an inner membrane that acts as a secondary active H^+^/drug antiporter (AdeB), extruding a vast spectrum of structurally unrelated drugs through a periplasmic membrane fusion protein channel (AdeA), connected to an outer membrane funnel shape channel (AdeC) (14, 15). The primary RND efflux pump of *A. baumannii* (AdeABC) was first described in MDR strain BM4454 (11). The *Ade*ABC locus has consisted of three tandemly linked genes encoding AdeA, AdeB, and AdeC proteins and showed resistance to several antibiotic classes such as aminoglycosides, tetracyclines, and fluoroquinolones, respectively (9). Expression of adeABC is tightly regulated by the two-component regulatory system AdeR-AdeS (40), encoded by the adeRS operon, located upstream from adeABC and transcribed in the opposite direction (16). The AdeB is an inner membrane protein and transports various drugs through the central pore opened to the periplasm via three vestibules located at subunit interfaces and a central cavity. More researchers demonstrated that the *ade*B is the most important gene in the adeABC efflux system (17,18). Cryo-Electron Microscopy (CEM) of the monomers of the inner membrane protein suggested that AdeB can adopt three different states: access (loose), binding (tight), and extrude (release) to provide essential dynamics for the efflux process (15). The AdeB trimer primarily adopts mainly a resting state (absence of antibiotics) with all promoters in a conformation devoid of transport channels or antibiotic binding sites (19).

Transmembrane proteins are relevant for drug development since they make up more than 50% of all human drug targets given that approximately 25% of all the proteins (20). Membrane proteins are difficult to study due to their partially hydrophobic surfaces, flexibility, and lack of stability (17). Consequently, this creates the need for bioinformatics tools that can accurately identify these types of proteins by predicting their topology, position, and orientation, as well as their N-terminal utilizing the contact map with coevolutionary precision matrices (21). Several methods have been developed over the last decades that predict protein topology with high accuracy. Many of these methods are freely available as web servers, both individually or combine with several other bioinformatics tools (22).

In the past 20 years, scientists were unable to produce effective antibiotics against Gram-positive or Gram-negative pathogens. Furthermore, the use of nanoparticles, gene therapy, phage therapy, and immunotherapy are either costly or still in their infancy (23, 24). In contrast, the amphiphilic antimicrobial peptides like Mastoparan B (a cationic, tetradecaeptide LKLKSIVSWAKKVL-CONH2 isolated from the venom of the hornet Vespa basalis) at low concentration has a potent antimicrobial activity and can be used along with antibiotics for treatment of various diseases (25). These compounds frequently destabilize biological membranes by dual amphiphilic activity resulting in membrane perforation (26, 27). In recent years, numerous efflux pump inhibitors have been discovered and tested, including natural products, antibiotics, and synthetic molecules (12). The combination of antibiotics with natural products circumvents the resistance phenomenon and consequently decreases the dose and drug side effects. The activity of these peptides in the combination with antibiotics on *Escherichia coli* AcrAB-TolC and the *Pseudomonas aeruginosa* MexAB-OprM complexes were well delineated (22). In contrast, there is no information about the effect of Mastoparan-B on the RND efflux in *A. baumannii*.

It has been reported that the phenylalanine-arginine β-naphthylamide (PAβN) a broad-spectrum efflux pump inhibitor interacts with AdeB strongly through the distal binding pocket (29). In another investigation, computational screening suggested that Strictamin (Akuammilan-17-oic acid methyl ester from Alstonia scholaris) and Limonin (7, 16-dioxo-7, 16-dideoxylimondiol from Citrus spp.) exhibited binding to RND efflux pump in MDR strains of *A. baumannii* (30).

Therefore, in the present study, for the first time, we are reporting transcriptional analysis of the *ade*B gene in the presence of Mastoparan-B, the detailed stereochemical structure of AdeB protein before and after binding to this peptide, superfamily analysis by the CATH database, Gene Ontology, molecular docking, prediction of structure conformation and helix folding by AlphaFold Protein Structure Database (AlphaFold 2). The emerging picture using this peptide in combination with antibiotics further suggests opportunities for drug designs.

## RESULTS

### Bacterial source and identification

In our previous reports, 65 isolates of *A. baumannii* were isolated from the ICUs hospitals in Kerman, Iran (31). Fifty-five of these were MDR, while 10 were carbapenems susceptible. Among them, one isolate that exhibited the highest MICs to various classes of antibiotics was selected for this study.

### Antimicrobial susceptibility tests and PCR-Sequencing

The organism exhibited high MICs against gentamicin and kanamycin (64 μg/ml), ciprofloxacin and levofloxacin (125 μg/ml), ceftazidime and ceftriaxone (256 μg/ml), imipenem (125 μg/ml), colistin (8 μg/ml), piperacillin/tazobactam (125 μg/ml), and azithromycin (64 μg/ml), respectively (Fig. 1A). Furthermore, MP-B prevented the active growth of the MDR strain of *A. baumannii* with mean MIC=1 μg/ml, where 99.9% of the cell’s population was not grown in Tryptic Soy broth medium (p≤0.05). A significant synergy was observed when the cells were grown in the presence of both antibiotics (various concentrations) and sub-MIC of MP-B (Fig. 1A). The MICs against gentamicin and kanamycin were reduced to 8 μg/ml, ciprofloxacin and levofloxacin to 4 μg/ml, ceftriaxone to 2 μg/ml, carbapenems to 4 μg/ml, and tetracycline to 2 μg/ml, respectively. However, no change in the MIC level of colistin, piperacillin-tazobactam, azithromycin, co-trimoxazole, and chloramphenicol was observed (Fig. 1A). To confirm the synergism of Mastoparan-B with antibiotics, we performed growth curve analysis of the isolate in the presence and absence of sub-MIC of MP-B. When cells grown in 0.5 μg/ml of MP-B, relatively steady growth was observed after 8 hours of incubation (Fig. 1B). The results indicated that the reduction in MIC against antibiotics was not due to the death of bacteria by Mastoparan-B but rather the inactivation of some cellular mechanism (s) such as efflux pump. The results were supported by plotting log10 CFU/ml survival at different concentrations of MP-B (Fig. 1C). As the figure indicates, the Mastoparan-B kills the bacteria and reduces the live population to zero within 2 h at high concentrations, while steady growth was observed at 0.5 μg/ml (sub-MIC level) concentration.

**Figure. 1.**
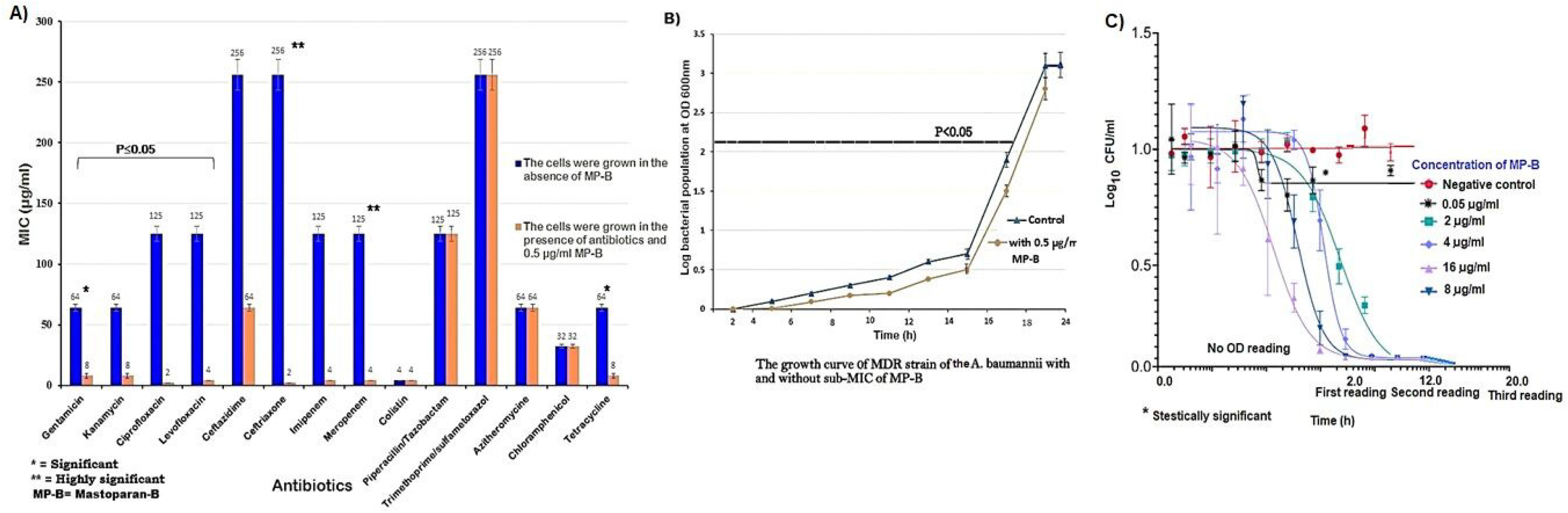
**A)** Antibiotic susceptibility of a multidrug-resistant *A. baumannii* isolates against 12 antibiotics in the presence and absence of sub-MIC concentration (0.5 μg/ml) of Mastoparan-B. The experiment was performed by broth microdilution test as described by CLSI -2019. **B)** Growth curve analysis of an *A. baumannii* isolate in the presence and absence of sub-MIC of MP-B. **C)** The survival rate of MDR *A. baumannii* in the presence of different concentrations of MP-B. The total cell death occurred within 2 h incubation in the presence of 16, 8, 4, and 1 μg/ml, while at 0.5 μg/ml concentration the growth is steady but lesser than the control. Values are presented as mean ± SD of three independent tests.

To prove this hypothesis, we performed the PCR technique by using specific primer pairs for the entire *ade*B gene. The AdeB efflux pump gene was then sequenced using Sanger’s dideoxy DNA chain termination method by ABI Prism DNA Sequencer, and its expression was evaluated in the presence and absence of MP-B. Interestingly, a 20-fold decrease in the expression of the *ade*B efflux pump gene was detected when the cells were grown at the sub-MIC level of this peptide. Furthermore, we obtained the nucleotide sequence of the *ade*B gene by blasting with similar sequences in the NCBI database (https://www.ncbi.nlm.nih.gov/blast). From the nucleotides, we detected the AdeB protein sequence in the GenBank database.

### Physicochemical parameters and Gene Ontology

The computation of physicochemical parameters of the AdeB efflux pump revealed that the protein was typically composed of 1036 amino acid residues with an average molecular weight (M. Wt.) of 112592.02 and the molar extinction coefficient of (ε) 81945 (Table 1A). Likewise, empirical analysis of amino acid composition by the Expasy ProtParam database revealed the dominance of small aliphatic amino acids like alanine at 16%, isoleucine, valine at 10%, and glycine at 6.7% in the AdeB structure (Table 1B). Furthermore, documentation resources of protein families, active domains, and functional sites indicated the AdeB protein belonged to the RND transporter efflux (HAE1) similar to ArcB protein Acrflvin-R, and ACR_tran family. It showed the nonhomologous relationship with superfamily ArcB_ DN_DC and pore-TolC-like domains. Gene Ontology further confirmed the biological, and functional activities of the AdeB protein. The overview of the sequence/structure diversity of the RND superfamily in this study compared to other superfamilies in the CATH database is presented in Fig. 2A. The obtained domains by CATH were structurally similar to MDR efflux transporter ArcB transmembrane domains with two related sequence families and one structural cluster (Fig. 2A). The AdeB efflux protein had 9.8% unique sequence annotations (Fig. 2B) and, 6 functional family members (lineage) in the *A. baumannii* (Fig. 2C).

**Table 1A.** Physicochemical parameters of AdeB efflux transporter from Acinetobacter baumannii used in this study.

| Parameter | Results |
| --- | --- |
| **Number of amino acids** | 1036 |
| **Molecular weight** | 112592.02 |
| **Theoretical pI** | 8.07 |
| **Atomic composition:** | Carbon C 5117<br>Hydrogen H 8212<br>Nitrogen N 1290<br>Oxygen O 1457<br>Sulfur S 46 |
| **Formula** | C5117H8212N1290O1457S46 |
| **Extinction coefficient** | 81945 |
| **Instability index** | 32.23 |
| **Aliphatic index** | 107.68 ± 0.02 |
| **GRAVY** | 0.351 |
| **MolProbity** | 0.79 |

**Table 1B:** The amino acids content of AdeB protein investigated in this study by Expasy-ProtParam tool.

| Amino acids | Percent (%) |
| --- | --- |
| Ala | 9.3 |
| Arg | 2.8 |
| Asn | 4.3 |
| Asp | 3.2 |
| Cys | 0.02 |
| Gln | 3.8 |
| Glu | 5.3 |
| Gly | 6.8 |
| His | 0.6 |
| Ile | 8.4 |
| Leu | 10.5 |
| Lys | 5.9 |
| Met | 4.2 |
| Phe | 5.4 |
| Pro | 4.4 |
| Ser | 8.2 |
| Thr | 5.5 |
| Trp | 1 |
| Tyr | 1.7 |
| Val | 8.5 |

Alanine was the most frequent residue in the protein followed by isoleucine and valine.

**Figure. 2.**
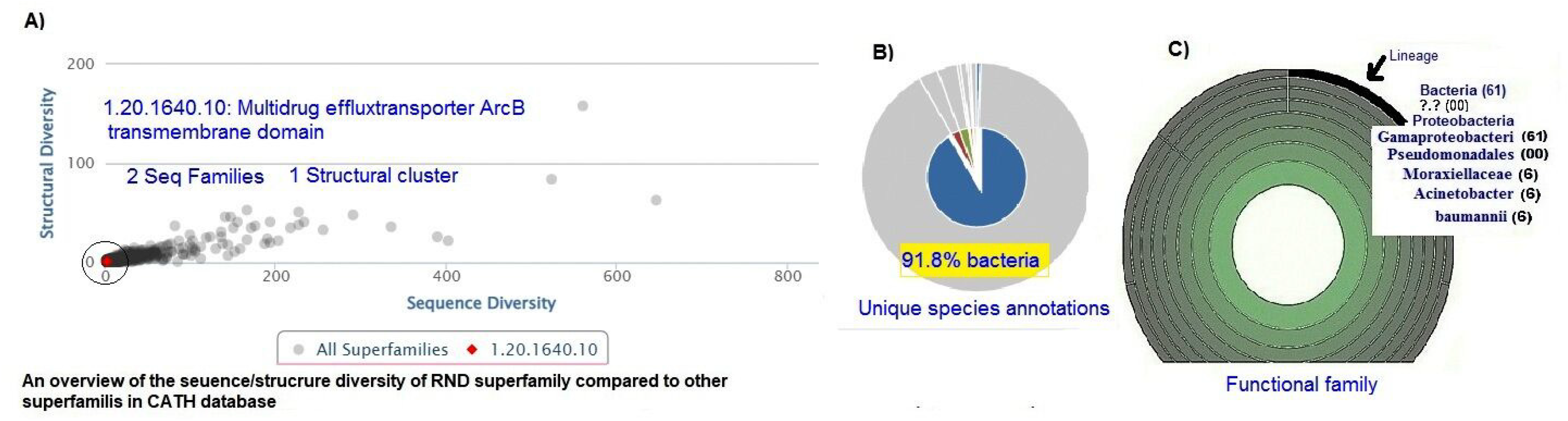
**A)** Overview of the sequence/structure diversity of the RND superfamily compared to other superfamilies in CATH (Class Architecture Topology Homologous superfamily), **B)** Unique species annotations and diversity, **C)** Six functional family in Morexiallaceae (*A. baumannii*), **D)** Superfamily superimposition with closely related CATH groups.

### Phylogenetic tree analysis and amino acids alignments

The phylogenetic tree analysis of the AdeB sequence convincingly demonstrated high sequence identity (99.9%) with AdeB efflux pump membrane transporter tr-AOA7L9E4V2_ACIBA, tr-A0A5P6FTF2_ACIBA, and tr**-**A0A3Q8ULEO_ACIBA, 99.6% sequence identity with tr-AOA373B8P5_9GAMM, and 99.41% identity with efflux pump tr-A0A086HUIO_ACIBA, respectively (Fig. 3A). Moreover, we investigated blast parameters including, identity (99%), score (Average 50300), and an E-value (5.3-11) (Fig. 3B). To prove the above results, we performed multiple amino acid alignments of the AdeB and compared the amino acids helix-5 for closely related strains, the results revealed high identity among the amino acids of these strains (Fig. 3C). Overall, the high identity matrix of the AdeB protein sequence was observed in the UniProtKB database (≤99%) (Fig. 3D). This indicates that a high evolutionary relationship exists among sequences in this report.

**Figure. 3.**
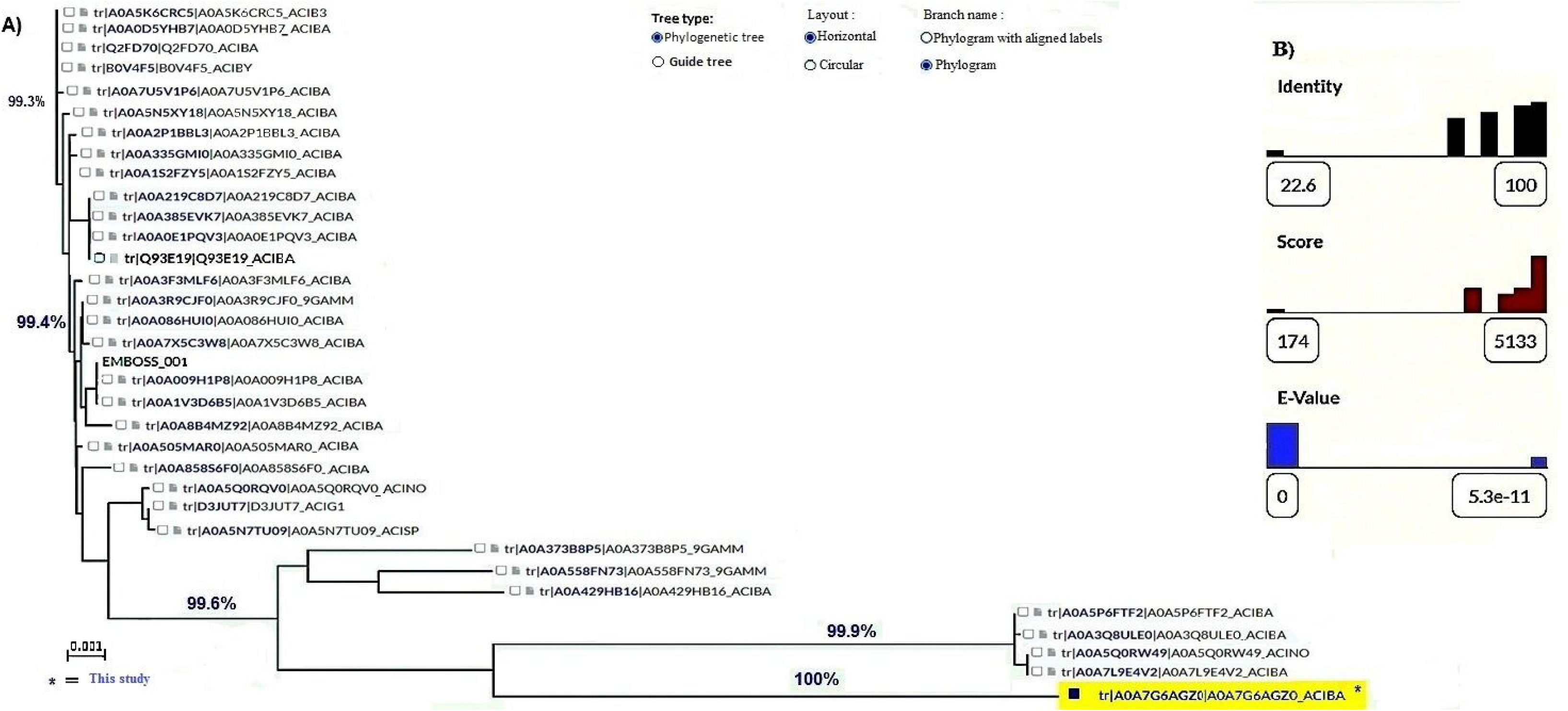
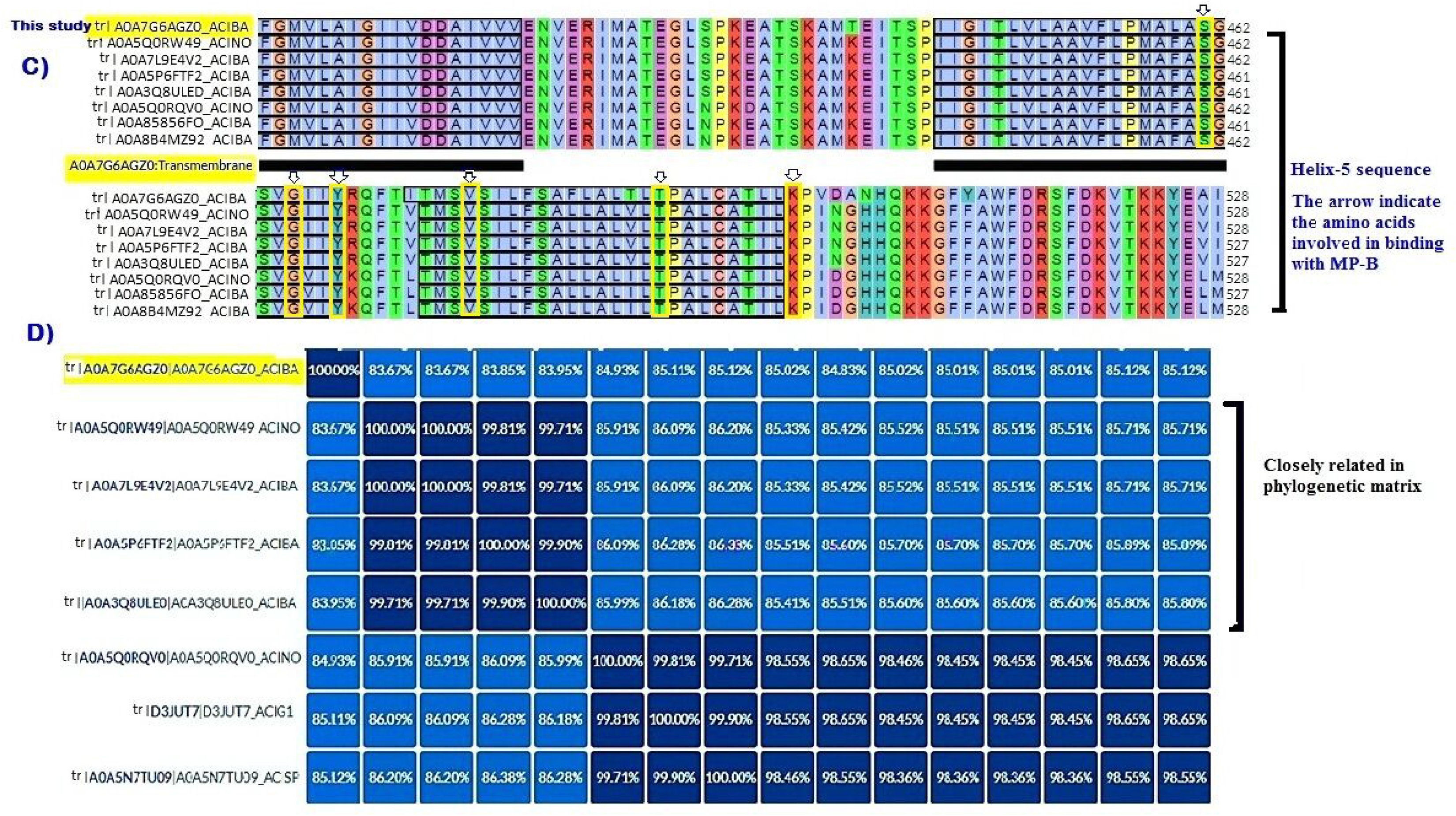
**A)** Phylogenetic tree evaluation of AdeB protein using UniProtKB database. **B)** AdeB sequence parameters including Identity, Score, and E-value. **C)** Multiple amino acid alignment of helix-5 of AdeB protein with closely related AdeB sequences in UniProtKB, **D)** Percent identity matrix among closest sequences (SD<99.4%).

### Molecular dynamics simulations of the efflux protein

The obtained AdeB trajectory showed an inner membrane protein with a 3-fold asymmetrical axis positioned perpendicular to the membrane surface with the typical RND-like folds. The predicted RND efflux pump exhibited three components; i) a 30Å U shape funnel channel in the AdeC outer membrane region composed of both α-helices and β-sheets (Fig. 4A), ii) a 40 Å large periplasmic membrane fusion AdeA region with the coil-coil conformation, and iii) a 50Å transmembrane AdeB transporter helical structure in the form of helix-turn-helix with a pore (Fig. 4A). The large periplasmic domain of the AdeB molecule is created by two extracellular loops that link together TM1 with TM2 and TM7 with TM8. This periplasmic domain can be divided into six subdomains, PN1, PN2, PC1, PC2, DN, and DC (Fig. 4B). These features fit with the extreme variety of antibiotics recognized by AdeB. The N-terminal TM1 leads to the PN1 subdomain that connects the PN2. Similarly, TM7 leads to the PC1 subdomain that was connected to the PC2 (Fig. 4B). Nevertheless, a pore connected the inner membrane cleft to the periplasmic cavity (Fig. 4C). A closer inspection of the pore reveals a high quantity of Lys, Phe, and Ala amino acids residues which relatively stabilized the pore channel (Fig. 4C). The combination of deep neural network learning with completed I-TASSER assembly significantly improved by threading template quality and therefore boosted the accuracy of the final model through optimized fragment assembly simulations with C-score = 1.41, TM-score =0.99, RMSD (Å) = 4.4 ± 2.9, and P-score = 1, respectively. Nevertheless, the validity of the AdeB structure was further confirmed by the Ramachandran plot (Fig. 5A) and local similarity estimate of amino acid residues in chains A, B, and C (Fig. 5B). The results indicated that over 99% of the amino acid backbones were in the Ramachandran favored regions and less in the outlier. The plot also illustrated torsional angles at low clash point Ψ (0.87%) along with high Ramachandran favored ϕ (97.92%) conditions. The obtained results of the local quality estimate by the Swiss-Expasy database for all three chains illustrated consistency in our work. Furthermore, the structural validation of the secondary protein structure using the PROSA website (Fig. 5C) showed high quality generated model with a Z-score = 2.3.

**Figure. 4.**
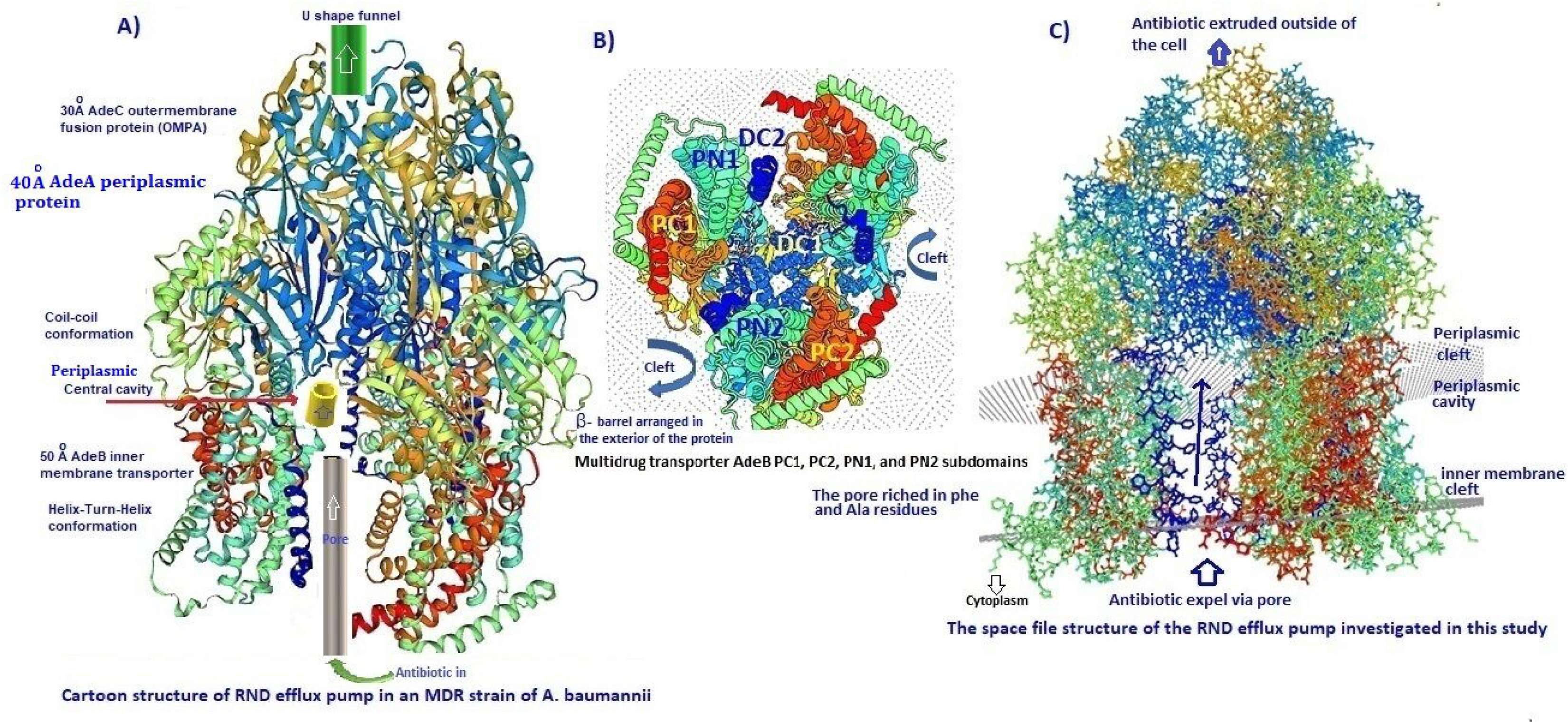
Schematic of representative structures of RND multidrug transporters tripartite assemblies. **A)** Cartoon structural prediction of the AdeABC RND efflux pump; a 30Å AdeC protein containing α/β barrel associated with a funnel and H^+^ antiporter fusion (OMPA) activity extrude antibiotic outside the cells. A 40Ǻ AdeA periplasmic protein with a heavy coil structure in the form of coil-coil conformation, the AdeA was associated with a large periplasmic cavity. A 50Ǻ the AdeB multidrug efflux pump inner membrane protein. The AdeB inner transmembrane helix contained a long narrow pore that connected to the central cavity. **B)** The distal end of AdeB contains PC1, PN1, PC2, and PN2 subdomains with three clefts. **C)** The wireframe diagram of the AdeABC RND efflux pump system shows the AdeB as an inner transmembrane helix, a narrow pore (rich in Phe, Lys, and Ala residues), a central cavity, a periplasmic fusion protein (OMPA), and 3 chains ABC associated with RND component.

**Figure. 5.**
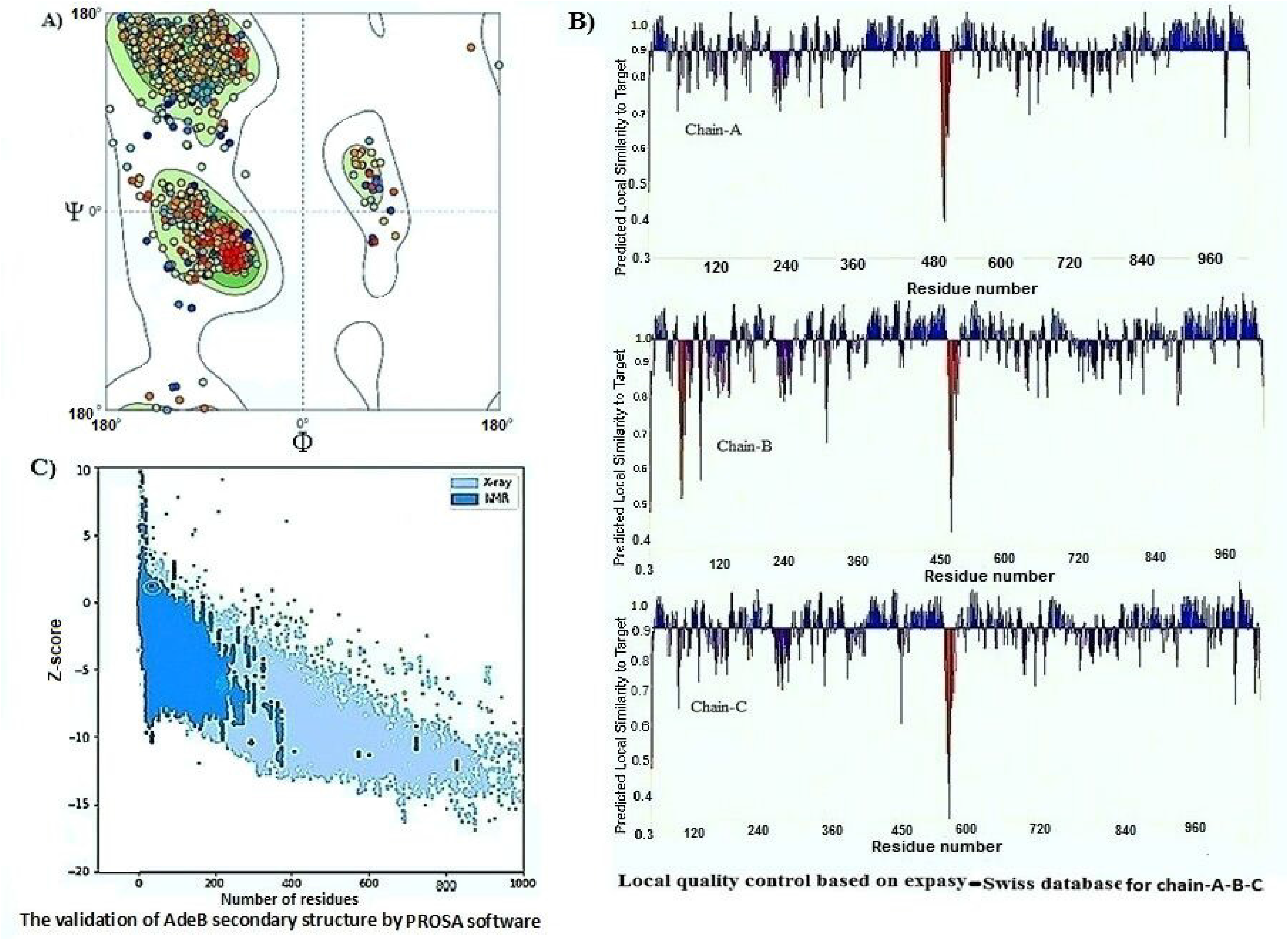
**A)** Ramachandran plot of AdeB protein based on homology modeling of Expasy database. The MolProbity score and clash points were estimated automatically. The majority of amino acids were in favor region (ε=0.79) which indicated energetically allowed regions for backbone dihedral angles Ψ and Φ of amino acid residues. **B)** Local quality estimate of the chains A, B, and C of AdeB protein using the Expasy program. **C)** ProSA Z-score of the AdeB efflux pump protein. The score of +2.12 indicated correct conformation of AdeB protein.

### Molecular docking analysis

For docking of Mastoparan-B with the AdeB protein, we prepared 20 poses and used them to determine the suitable binding site on the AdeB efflux pump. The comparison of the molecular docking simulation before and after attachment of MP-B showed MP-B attached exclusively with the helix-5 inner membrane via Val 499, Phe 454, Thr 474, Ser 461, Gly 465, and Tyr 468 residues, respectively (Fig. 6A). Upon docking of the ligand, there was a change in energy minimization (glide score of 0.8 to 0.2), and a shift in the orientation of dihedral angles including phi (φ), and psi (Ψ) by distances of 9.0 Å, 9.3 Å, and 9.6 Å (Fig. 6B). The black arrows indicate the bond interaction potentials: bond-stretching, VB, angle-bending, VA, dihedral (out-of-plane), Vdh. The protein folding changes were detected by AlphaFold DB software. Here, we used the Alfafold database for the first time for the determination of the AdeB domains and detailed information about folding and the position of helix-5 in the interior side of the AdeB protein (Fig. 7A). In addition, the attachment of MP-B to this helix and amino acids is involved in molecular docking studied by this software shown in Fig.7B. The structure was prepared, at physiological pH, using the Protein Preparation wizard in Schr ö dinger’s Maestro. We found a key hydrogen bonding interaction between the heteroatoms of Ala, Thr, and Gly plays an important role in binding to the MP-B α-helix structure. The overall docking process, positions of the amino acids, and H-bonds with the ligand molecule and receptor are shown in Fig. 7C. Moreover, comparing the AdeB molecule with template 7cz9A as the model indicated a high score in the ligand-receptor interaction (z-score = 33.510 cutoff = 6.9).

**Figure. 6.**
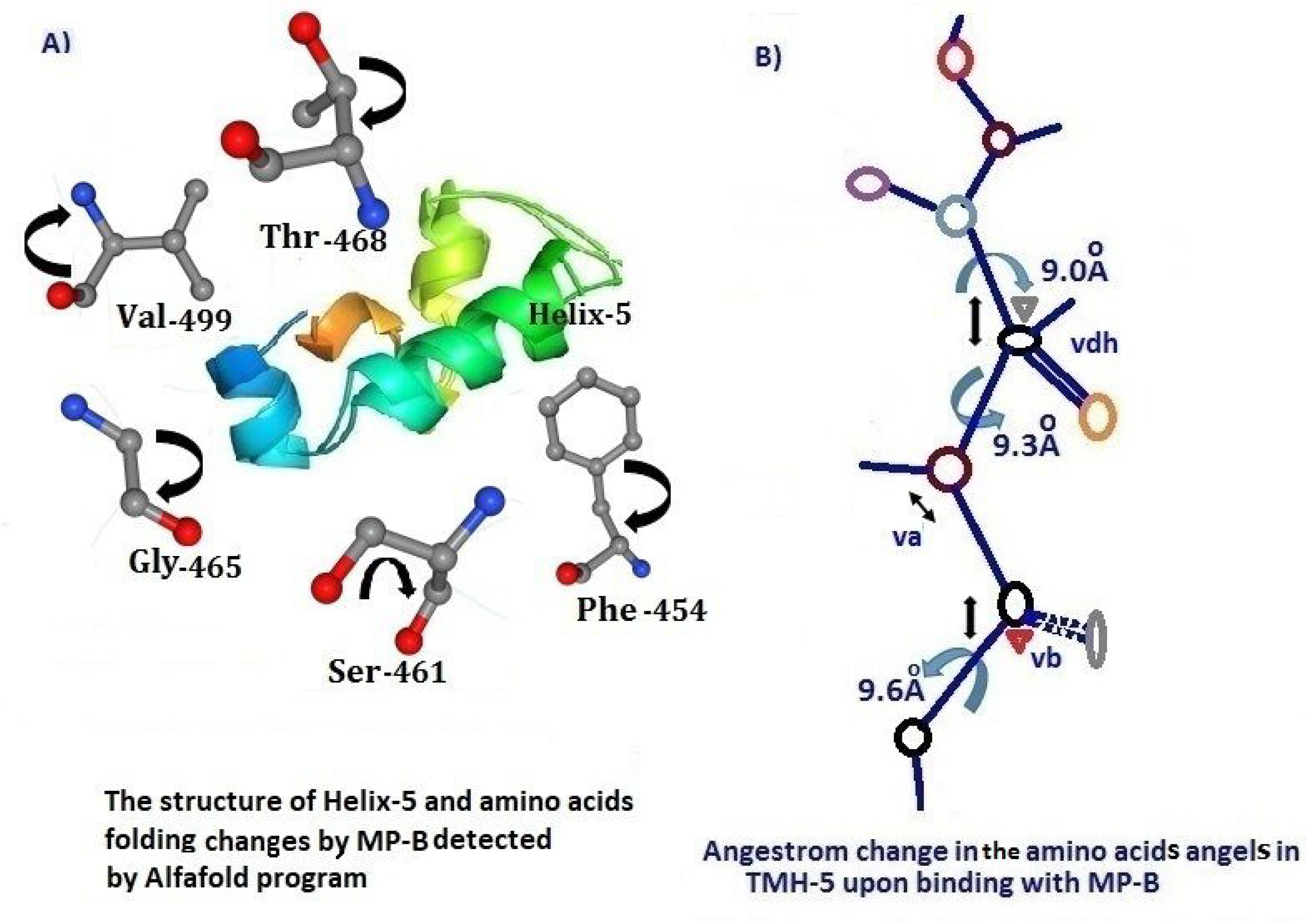
**A)** The folding of transmembrane helix-5 and angstrom changes in the amino acids involved in binding with MP-B (black arrow). The correct folding at atomic level obtained by AlphaFold 2 program, **B)** A graphical representation of the shift in amino acids angles (Ψ/ϕ) by 9.0 Å, 9.3 Å, and 9.6 Å.

**Figure. 7.**
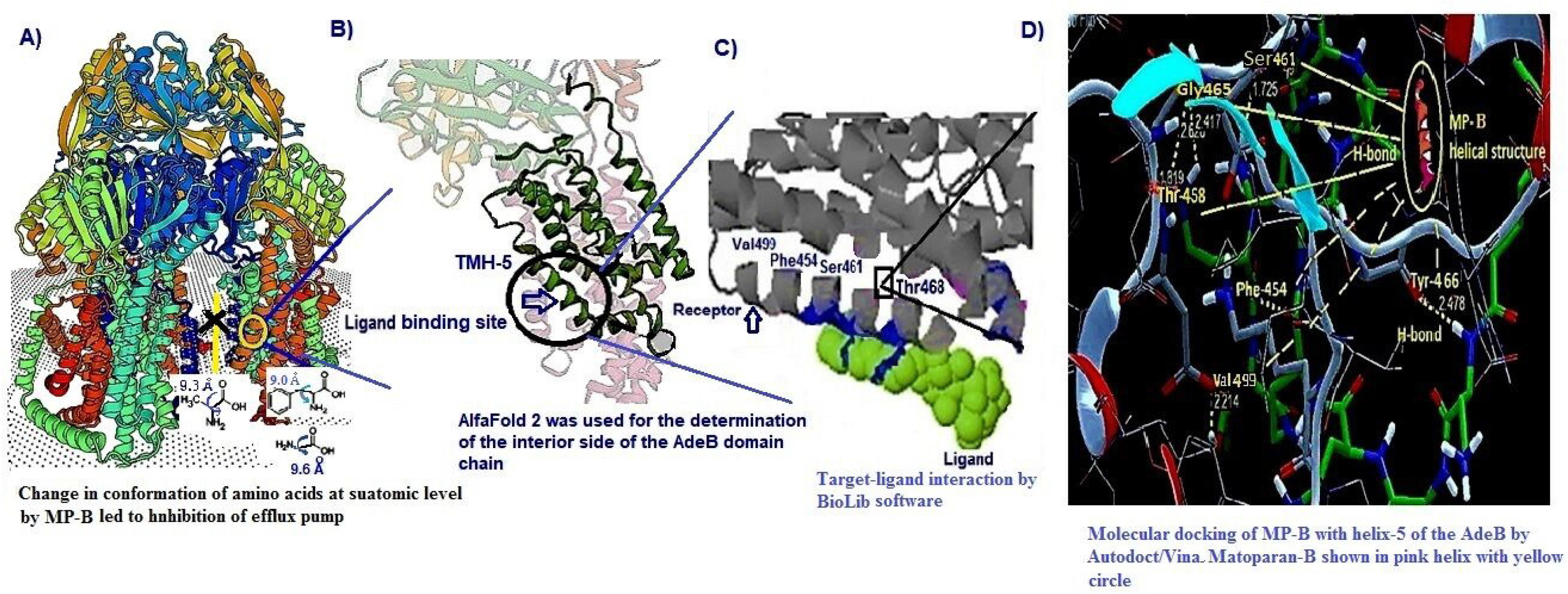
**A)** Schematic illustration of the AdeB binding site and internal domain chain depicted by AlphaFold 2 program, **B)** The illustration of AdeB domain chain including transmembrane helix-5, **C)** Amino acids in the helix-5 involved in molecular docking with Mastoparan-B, **D)** Binding of the best pose of α-helix to MP-B (The ligand showed by a yellow color small circle), **E)** Virtual position of amino acids form H-bounds attachment with MP-B via using AutoDock/Vina. The amino acids involved in docking are shown in yellow color.

### Prediction of position and topology of transmembrane helices

The topology of AdeB protein was predicted by both DeepTMHMM and TOPCONS software using Z-coordinates by prediction of ΔG-values across the sequence and Wilcoxon rank-sum test to get a quantified comparison between the two reliability scores independent of the overall prediction accuracy. The majority of transmembrane helices were arranged at positions between 300-500 and 900-1000. The place of binding ligand to the receptor is shown by a circle with a Z score of 5.8 (Fig. 8A). Consequently, analysis by TOPOCONS further confirmed the above estimate (Fig. 8B), and showed the percent of α-helix inside, membrane, and outside of the AdeB protein (Fig. 8C). The transmembrane helices (TMH) topology constituted the highest percentage in the AdeB structure. Furthermore, by employing Pyr^2^ software, we determined 47.7% of amino acid residues displayed α-helical structure, 21% β-sheets, 11.5% turns, and 24.5% coil with the Z score +2.52. As a result, the most frequent topology in AdeB protein was α-helix, followed by a random coil and extended β-strand barrel.

**Figure. 8.**
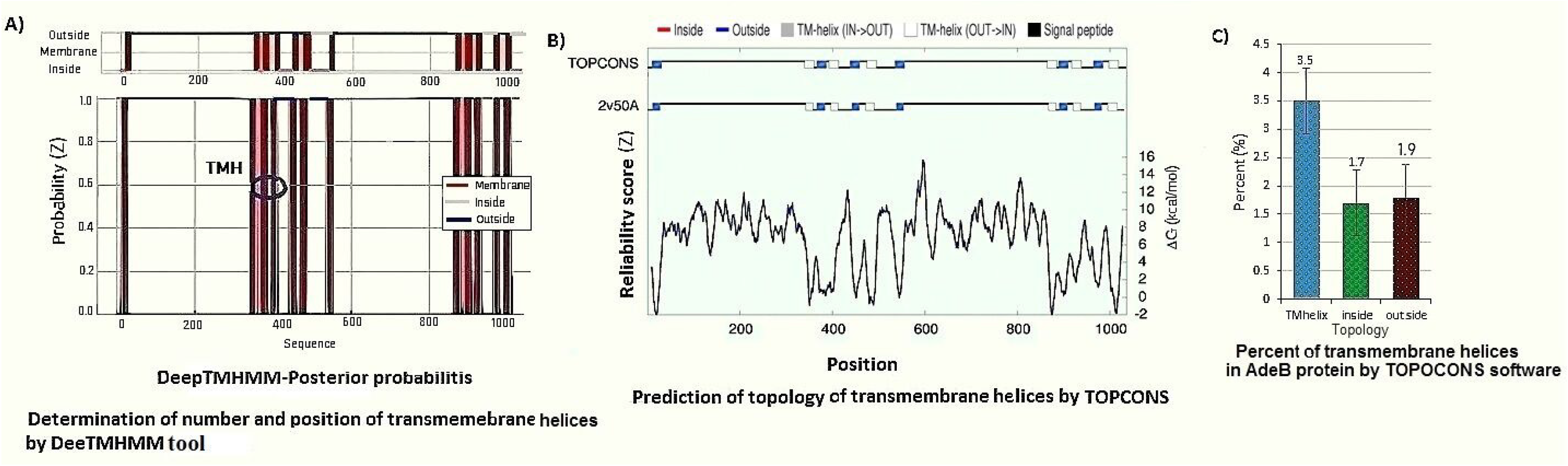
**A)** Determination of the number and position of transmembrane helices by the DeepTMHMM program (http://dtu.biolib.com/DeepTMHMM/), **B)** Prediction of the AdeB topology by the TOPOCONS software (https://topcons.cbr.su.se/), **C)** Percent of, inside, transmembrane, and outside helices using the TOPOCONS tool.

### Contact maps of the AdeB protein

The contact map analysis by the FUpred server represents a range of 1-20 angstroms. There are three overlapping D1-3, D6-2 and D6-1 at the N-terminal segment with a confidence score of 8.76 (Fig. 9A). The contact maps derived from the application of a distance cutoff of 9 to 11Å around the Cβ atoms constitute the most accurate representation of the visualized 3D structure. Nevertheless, the domain boundaries were shown by a continuous domain curve (Fig. 9B). This was followed by heat map scores which showed a low domain shift point for AdeB protein (Fig. 9C). Overall, the predicted continuous three-domain protein secondary structure of the AdeB suggested a confidence score of >8.6, indicating our prediction was accurate.

**Figure. 9.**
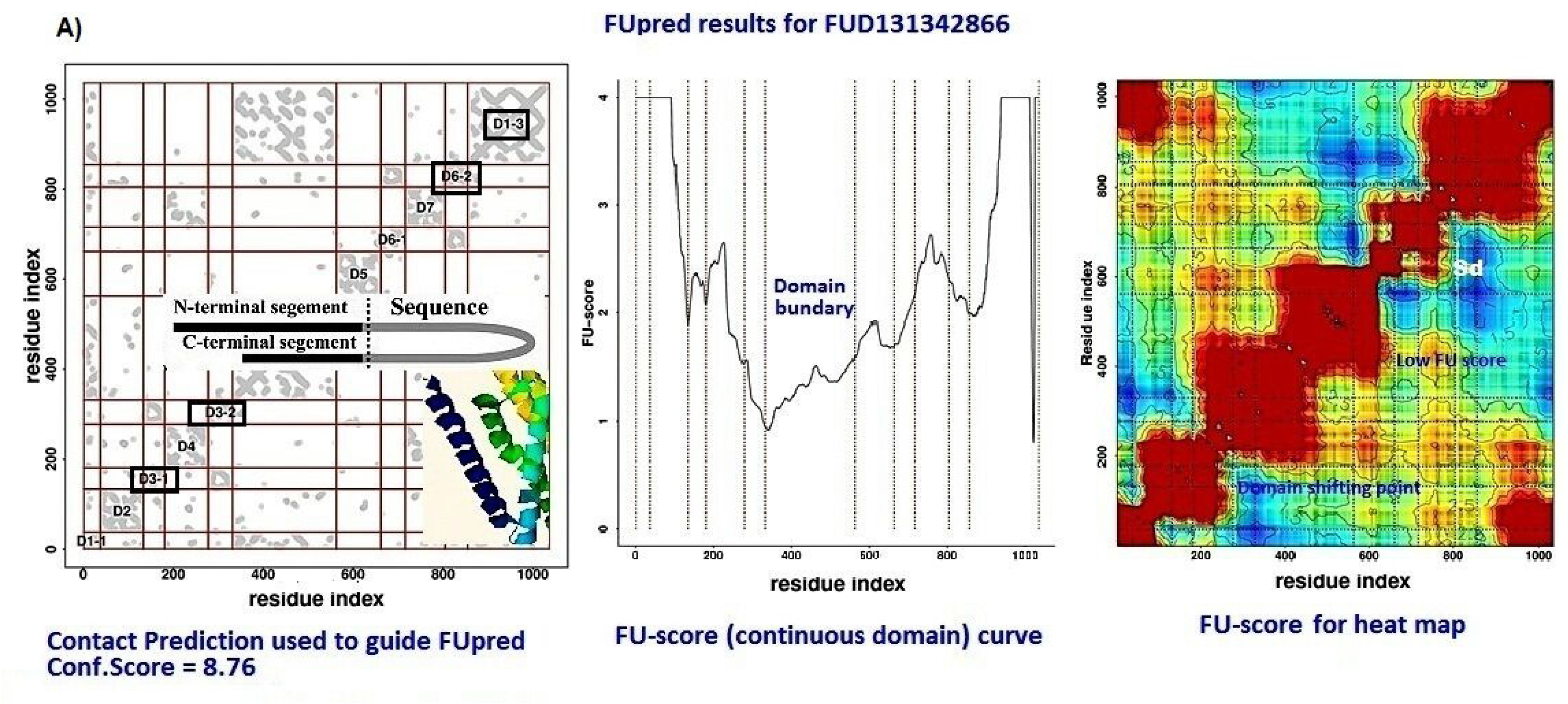
**A)** Contact map measured by FUpred software for accuracy of domains, **B)** Fu score for domains boundaries. **C)** FU score for heat map which shows domain shifting point of the AdeB efflux pump protein. The axes marked the residue index along the sequence in the contact map. For the contact map, each dot represents a residue pair with predicted contact. The terminal contacts and the corresponding inter-chain interaction are highlighted with a black box.

## DISCUSSION

A key part of the defense of *A. baumannii* against different antibiotics is a series of efflux pumps, which effectively extrude a large number of antibiotics, making the pathogen intrinsically more resistant against antibiotics (32). The efflux pumps are a topic of considerable interest, both from the perspective of understanding function and as targets for novel adjunct therapies (33). In *A. baumannii*, three RND efflux systems, AdeABC, AdeFGH, and AdeIJK are reported to be primarily associated with the emergence of MDR strains (31). Nevertheless, the RND efflux family in *A. baumannii* has broad substrate specificity and can efflux a vast variety of antibiotics from the periplasm before the antibiotics even fully enter the cell (34).

Despite a large number of identified efflux pumps, our understanding of the molecular mechanisms by which the antimicrobial drugs remain limited. Although relationships between drugs and targets have been depicted in a global view, the relationship between amphiphilic peptides and the AdeB efflux pump remains uncharacterized. Moreover, most of the current drug targets are proteins, and the 3D structures of these proteins need to be known in target identification. Therefore, computer-aided drug target identification methods can greatly reduce the searching scope of experimental targets and associated costs by identifying the d-binding sites and evaluating the druggability of the predicted active sites (35).

The results from this study provide novel insight into the inhibitory activity of Mastoparan-B on the RND efflux pump. Our target was AdeB protein in MDR *A. baumannii*. The preliminary susceptibility test revealed that the organism was intrinsically sensitive to MP-B as a potent inhibitor of the membrane function. However, when the cells were grown in the presence of a sub-MIC concentration of this peptide, and different concentrations of antibiotics, there was a significant reduction in MICs particularly aminoglycosides, fluoroquinolones, third-generation of cephalosporins, carbapenems, and tetracycline, respectively. In contrast, the MIC against colistin, piperacillin-tazobactam, azithromycin, trimethoprim-sulfamethoxazole, and chloramphenicol remained unchanged. This led us to the understanding that, the AdeB efflux pump might be involved in this extrusion of particular antibiotics in its channel. The synergic activity of MP-B with antibiotics was further supported by the transcriptional analysis of the *ade*B gene in the presence of MP-B which encode the AdeB protein. Furthermore, the search for phylogenetic tree analysis of our AdeB protein with similar sequences in the UniProtKB database revealed a low percent identity matrix using the Maximum Likelihood method. The results indicated the AdeB investigated in this study was not evolutionary-related to the majority of Acinetobacter efflux pumps.

Furthermore, we performed docking and free energy change, revealing that Mastoparan-B with amphiphilic α-helical structure binds to the inner membrane AdeB protein via the hydrophobic amino acids. These nonpolar amino acids have either aliphatic or aromatic side chains. By AlphaFold tool, we found that the proteins will fold into a 3D dimensional shape to bury these hydrophobic side chains in the protein interior. Since there was no reliable crystallography for a membrane protein, alternatively, one can resort to computational methods. These methods are based on structural, physicochemical, and evolutionary properties that distinguish binding sites from the rest of the protein surface (e.g., amino acid composition and residue conservation). Furthermore, we investigate deep learning of artificial intelligence for the prediction of the membrane topology of transmembrane proteins. A critical step in the biosynthetic pathway leading to the folded protein in the membrane is its insertion into the lipid bilayer. Understanding the fundamentals of the insertion and folding processes will significantly improve the methods used to predict the three-dimensional membrane protein structure from the amino acid sequence (36). The attachment of all forms of MP-B exclusively occurred to the transmembrane helix-5 which is situated at the interior of the inner membrane domain. This was confirmed by using the BioLiP database (the structure was predicted by SPARKS-X (SPX) with template 7cz9A. z-score=33.510 (cutoff=6.9). Concurrent target-ligand interaction strongly indicates a shift in the dihedral angles with a lower free binding energy score. Consequently, this phenomenon caused a transient conformational change in the protein structure resulting in the inhibition of AdeB activity. Similarly, the presence of a large amount of Phe, Ala, and Lys amino acid residues in the pore region provides a hydrophobic trap around this peptide in the AdeB molecule. Particularly, Phe residue was important in creating this hydrophobic microenvironment. So far, the mechanisms have not been studied in detail, therefore this statement must be further confirmed. Similarly, we used FUpred, which predicts protein domain boundaries utilizing contact maps created by deep residual neural networks coupled with coevolutionary precision matrices. The results indicated several interchain transfers at C-terminal like D1-3, D3-2, and D6-2 discontinuous three-domain protein using the following software https://zhanglab.ccmb.med.umich.edu/FUpred.

## MATERIALS AND METHODS

### Bacterial origin and PCR-Sequencing

An MDR strain of *A. baumannii* was isolated from the previous collection (31) and used for the detection of the *ade*B gene by polymerase chain reaction (PCR) technique. Briefly, total genomic DNA was extracted by a DNA genomic extraction kit (ParsTous, Iran). The amplification was carried out using a common PCR core mixture (total volume, 50 μl) containing 20 μL PCR buffer, 10 μL of 200 mM each dNTP (Bioneer, South Korea), 1 U of AmpliTaq DNA polymerase (ParsTous, Iran), 20 pmol of the forward -5’ TTAACGATAGCGTTGTAACC -3’ and reverse 5’-TGAGCAGACAATGGAATAGT -3’ primer pair (Bioneer, Korea) and 50 ng of the genomic DNA as the PCR template. The *ade*B gene was further purified by a DNA purification kit (ParsTous, Iran) and sequenced in both directions by Sanger’s dideoxy chain termination method using ABI Prism 373 DNA Sequencer (Applied Biosystems 373/3730XL, Bioneer, Korea). The *ade*B sequence was then submitted to the GenBank NCBI database (https://www.ncbi.nlm.nih.gov/genbank/submit/) for accession number. The BLAST program was used to find the amino acid sequence of the AdeB protein.

### Susceptibility to antibiotics and Mastoparan-B

The susceptibility of the selected strain against 12 antibiotics was carried out by disc diffusion assay and MIC methods using broth microdilution test as suggested by the Clinical and Laboratory Standards Institute (CLSI 2019) (33). At the same time, the susceptibility of the isolate to Mastoparan-B was performed by microdilution method using the Tryptic Soy broth medium, and criteria for sensitivity or resistance were measured based on our previous report (38). Similarly, we performed MICs of antibiotics in the presence of sub-MIC concentration (0.5 μg/ml) of MP-B to see any synergism between peptide and antibiotics. Before the experiment, we checked the viability of the cells at this concentration, by removing 1 ml aliquots of each sample grown in the presence and absence of 0.5 ml of MP-B at each 2 h and reading the turbidity with a UV/Vis spectrophotometer (Persia Med, AE-S60-4U) at an optical density (OD) of 600 nm. The standard cultures of *E. coli* ATTC25922 and *A. baumannii* ATCC 19606 were used as the quality control strains. All antibiotics were purchased from MAST Company, Manchester, UK. Mastoparan-B powder with 99.5% purity was purchased from Shimi-Daru (Tehran, Iran) and used as described by the manufacturer.

### Expressional analyses of the adeB gene

RNA extraction, C-DNA synthesis, and relative quantitative real-time PCR (qRT-PCR) were performed to assess the expression of the *ade*B gene in the presence and absence of 0.5 μg/ml of MP-B as described previously (31). Briefly, amplification was done using 25 μl of SYBR®Green dye and 2× Real-Time PCR master mix solution (BIO FACT, Korea), followed by the addition of 1 μl of *ade*B primer pair F: 5’-AACGGACGACCATCTTTGAGTATT-3’ and R: 5-’CAGTTGTTCCATTTCACGCATT-3’ and 5 μl of cDNA template as described by the manufacturer. Quantification of the *ade*B gene was performed by using the ABI Step One qRT-PCR system (Applied Biosystems, Foster City, CA, USA). The gene expression was calculated as a fold change (RQ) between the target gene and matched reference 16SrRNA levels based on the following formula RQ = ^2Δ^DDCt. Where ΔCt is equal to the difference between the cutting point (Ct) value for the analyzed gene and Ct for the 16SrRNA reference gene as described by Livak et al. (39). *A. baumannii* ATCC 19660 was used as the internal control strain. Differences were assessed by Student’s t-test for consideration as statistically significant.

### Classification, physicochemical parameters, and Gene Ontology

The hierarchical classification of AdeB protein to the superfamily, homologous family, predicting domains, and important functional sites were carried out by the InterPro software suite (https://www.ebi.ac.uk/interpro/). To classify proteins in this way, InterPro uses predictive models, known as signatures provided by several different alignments, and produces a comprehensive resource for protein classification (https://www.ebi.ac.uk/interpro/search/sequence/). Any alignment with a TM-score ≥0.5, which is the cutoff value for a significant structural match, was used as a potential template for the target protein-peptide interaction. Furthermore, molecular weight, theoretical pI, amino acid composition, atomic composition, extinction coefficient, estimated half-life, instability index, aliphatic index, and grand average of hydropathicity (GRAVY) were computed by information stored in the Swiss-Port ProtParam database (https://web.expasy.org/protparam/). Gene Ontology at molecular and cellular functions was determined using the UniProtKB platform (https://www.uniprot.org/). We also worked on the domain-based method for the AdeB protein sequence versus structural diversity and prediction of the functional site that exploits functional sub-classification of the superfamilies by the CATH protein classification database (https://www.cathdb.info/).

### Construction of phylogenetic tree of the AdeB protein

The phylogenetic tree of the AdeB protein was constructed using maximum likelihood sequence alignment Clustral Omega version 7 in the UniProtKB pipeline, and variable sites were removed by the stringent settings (40). A bootstrapped was generated with RAxML (Randomized Accelerated Maximum Likelihood) version 7.0.3 with the confidence levels (%) generated from 1000 bootstrap trials. Moreover, we choose a consensus sequence manually using similar protein sequences from the UniProtKB restricted to the *A. baumannii* AdeB protein.

### Homology modeling and prediction of protein folding by the AlphaFold 2 database

From the primary amino acid sequence, the 3D model of AdeB protein was generated by the SWISS-Expasy (https://swissmodel.expasy.org), and I-TESSAR (https://zhanggroup.org/I-TASSER/) platforms using the homology modeling program. For each target, I-TASSER simulation generates a large ensemble of structural conformations, called decoys. To select the final models, I-TASSER uses the SPICKER program to cluster all the decoys based on the pair-wise structural similarity and reports up the top five models which correspond to the five closest structure clusters. The confidence of each model is quantitatively measured by the C-score that is calculated based on the significance of threading template alignments and the convergence parameters of the structure assembly simulations. Furthermore, for the evaluation of multiple, amino acids alignments and similarity matrix, we used the FASTA format of the targeted amino acids sequence and blasted in the UniProtKB database (https://www.uniprot.org/). The functional domains of the AdeB protein were bolded using the ROSETTA prediction tool (https://www.rosettacommons.org/) and the Hidden Markov Model program (HMMs) (https://www.ebi.ac.uk/Tools/hmmer/). The quality of obtained models was validated with a comprehensive scoring function for model quality assessment servers such as ProSA and QMEAN (41, 42). The dihedral angles including phi (φ) and psi (Ψ) and backbone conformation were assessed using the PROCHECK tool at the PDB Sum server (https://www.ebi.ac.uk/thornton-srv/software/PROCHECK/). In this study, we evaluated the accuracy of the AdeB protein structure, folding the helices and domain side chains including the interior side of the AdeB by the AlphaFold 2 database (https://alphafold.ebi.ac.uk). In addition, the backbone dihedral angles against amino acid residues and amino acid clash points were determined by the Ramachandran plot (44). The final ensemble of structures is well-defined with an average pairwise root-mean-square deviation (RMSD) across backbone atoms of 0.71 Å and an average MolProbity score of 1.8.

### Prediction of the protein-peptide interaction site

Because of peptide flexibility and the transient nature of protein-peptide interactions, peptides are difficult to study experimentally. Therefore, we used computational methods for predicting structural information about protein-peptide interactions by the InterPep server (http://wallnerlab.org/InterPep/). The InterPep software is a powerful tool for identifying peptide-binding sites with a precision of 80% at a recall, of 20%. It is an excellent starting point for docking protocols or experiments investigating peptide interactions.

### Ligand preparation and molecular docking

Mastoparan-B with MF: C78H138N20O16 was retrieved from the PubChem database (Compound CID: 86289587) and used as a ligand in this study. 20 different ligand poses were selected using the LigPrep module in Maestro v11 employing the Schrödinger platform (44). The best interaction poses were visualized using the PyMOL server (https://pymol.org). The most comprehensive and accurate model of ligand-protein docking and the protein function annotation was obtained from the BioLiP database (http://zhanglab.ccmb.med.umich.edu/BioLiP/). In brief, the free energy was minimized by the Universal Force Field and converted to the pdbqt format in PyRx0.8 for virtual screening (45) Furthermore, the favorable potential drug target interactions between the selected ligand molecule and modeled receptor (AdeB) were identified by the extra precision (XP) feature of Grid-based Ligand Docking with Energetics (GLIDE) in the Schrödinger platform (44). Each simulation was repeated three times with different initial conditions to increase the precision of the simulations and to prevent any dependencies of the results on the initial conditions. Binding sites were generated using the Site Map tool version 2.4 (Schrödinger Inc.). Finally, molecular docking was performed via AutoDock/Vina suite to assign the ligands and receptors bond orders by adding hydrogen atoms (46). For docking purposes, the best docking pose was selected directly using the Glide G-Score. Once G-Score was obtained, it was used to build optimal binding free energy that distinguishes our sequence from the reference sequence (usually taken to be the optimal sequence). Therefore, Site Map was tasked to identify the five top-ranked possible receptor suites using the default settings which include size, volume, amino acid exposure, enclosure, contact, hydrophobicity, hydrophilicity, and donor/acceptor ratio. Furthermore, the topology of the transmembrane, inside and outside α-helices was predicted by the DeepTMHMM (https://dtu.biolib.com/DeepTMHMM) and TOPCONS (http://topcons.net/) modules.

### Preparation of the contact map and domain boundaries

We predicted the AdeB protein contact map and domain boundaries using deep residual neural networks coupled with coevolutionary precision matrices (https://zhanglab.ccmb.med.umich.edu/FUpred). A protein contact map represents the distance between all possible amino acid residue pairs of a 3D structure. For two residues, the element of the matrix is 1 if the two residues are closer than a predetermined threshold, and 0 otherwise (47). To achieve reliable results, we used the FUpred scoring software for continuous two-domain protein, using the following formula

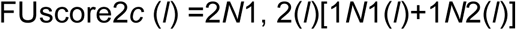

where *l* is the domain splitting point of a protein, *N*_1_ (*l*) and *N*_2_ (*l*) represent the number of contacts within the first and second domains, respectively, and *N*_1, 2_(*l*) = *N*_2, 1_(*l*) indicates the number of contacts between the first and second domains.^48^ The core idea of the algorithm was to retrieve domains boundaries locations by maximizing the number of intradomain contacts while minimizing the number of interdomain contacts from the contact map.

## Statistical analysis

Statistical analyses were performed using SPSS 17.0 (SPSS, Chicago, IL, USA). A p ≤ 0.05 was considered statistically significant for two-tailed tests.

## Conclusion

AdeB is a critically important representative of the RND family of multidrug transporters in *A. baumannii* and as such, the molecular interaction of this pump with Mastoparan-B by bioinformatics tools will provide useful new information on the targeted inhibition of efflux pumps in this bacterium. Mastoparan-B might be used in combination with antibiotics like ampicillin-sulbactam for the treatment of MDR *A. baumannii*.

## Conflicting interests

The authors declare that they have no competing interests.

## Data availability

The authors confirm that the data supporting the findings of this study are available within the article and its supplementary materials

## Authors’ contributions

S.MR performed the analysis and participated in the design of the study, OZ, and OT, helped in writing and correcting of paper, FM helped drafted the manuscript and analysis of data, and G.MM help with software collection and information analysis. All authors read and approved the final manuscript

## Funding

Not applicable.

